# Wireless monitoring of respiration with EEG reveals relationships between respiration, behaviour and brain activity in freely moving mice

**DOI:** 10.1101/2023.06.19.544346

**Authors:** Debanjan Dasgupta, Deborah Schneider-Luftman, Andreas Schaefer, Julia J. Harris

**Author notes:** These authors contributed equally (Debanjan Dasgupta, Julia J. Harris).

## Abstract

Active sampling in the olfactory domain is an important aspect of mouse behaviour. Numerous methods are being used to measure active sampling behaviour, yet reliable observation of respiration in untethered, freely moving animals is challenging. So far, methods for measuring this have largely been restricted to head-fixed sniff monitoring, which makes it difficult to understand how sniff changes are related to natural mouse behaviour. Here, we implant a telemetry-based pressure sensor into the right jugular vein, which allows respiration to be measured via wireless thoracic pressure sensing in awake and freely moving, untethered mice. After verifying this technique against standard head-fixed respiration measurements, we investigated respiration patterns across a range of experiments in freely moving animals. Respiration frequency increased as mice voluntarily explored novel environmental cues. Combining wireless respiration measurements with EEG/EMG recording, we then used an evolving partial coherence analysis to uncover the direct relationships between respiration and brain activity in different frequency bands over the same exploration period. Finally, we examined respiration patterns across different vigilance states, revealing changes in passive respiration frequency across wakefulness, deep (NREM) sleep and dreaming (REM) sleep, and odour-triggered respiration increases in the absence of brain activity changes during NREM sleep. As it can be combined with behavioural assays and brain recordings, we anticipate that wireless respiration monitoring will be a valuable tool to increase our understanding of how mice use olfaction to process and interact with the environment around them.

## INTRODUCTION

Active sampling plays a crucial role in sensory processing (Schroeder at al., 2010), particularly within the sense of olfaction (Margrie & Schaefer, 2003; Kepecs et al., 2006, Verhagen et al., 2007; Cenier et al., 2013; Jordan et al., 2018a,b; Shusterman et al., 2018). Olfactory sampling is governed by the respiration rhythm, and many mammals display a huge repertoire of “sniffing” behaviours which dynamically alter their respiration rate (Welker, 1964; Youngentob, 1987; Wachowiak, 2011; Jordan et al., 2018a,b). In turn, respiration-entrained neuronal firing patterns are increasingly being observed across brain regions outside of the olfactory system, including the hippocampus, neocortex and limbic system (reviewed in Folschweiller & Sauer, 2023). Recent evidence has led to the hypothesis that breathing may set a global brain rhythm to actively coordinate neuronal communication across distant brain regions (Heck et al., 2017; Tort et al., 2018), and it is now being discovered that the alignment of different neural activity patterns to this respiration rhythm can change with the arousal state of the animal (Zhong et al., 2017; Cavelli et al., 2020; Girin et al., 2021; Tort et al., 2021; Karalis & Sirota, 2022). During exploration, respiratory rhythms overlap in frequency with the hippocampal theta rhythm (Nguyen Chi et al., 2016) and confusion may therefore arise in the absence of knowledge about the animal’s breathing. Accurately measuring sniffing is therefore critical not only in the fields of respiratory physiology and olfaction, but also in the context of animal behaviour and neural processing more generally.

Over the years, different strategies have been developed to monitor respiration (Grimaud & Murthy, 2018), including whole-body plethysmography (Walker et al., 1997; Matrot et al., 2005; Lim et al., 2014; Prada-Dacasa et al., 2020); monitoring chest distension using piezoelectric bands (Fukunaga et al., 2012; Zehendner et al., 2013); thermal imaging of nostrils (Esquivelzeta Rabell et al., 2017); and monitoring changes in nasal air flow (Oka et al., 2009; Fukunaga et al., 2012; Bolding and Franks, 2017; Ackels et al., 2021; Dasgupta et al., 2022), pressure (Shusterman et al., 2011; Smear et al., 2011; Reisert et al., 2014; Sirotin et al., 2014; Jordan et al., 2018), or temperature (Macrides, 1975; Ito et al., 2014; McAfee et al., 2016; Liu and Han, 2022; Liu et al., 2022). However, these methods all require specific recording conditions or tethering, making it difficult to measure respiration during natural behaviours or in combination with other recording techniques. Recently, an innovative approach was developed, which has the potential to overcome this challenge. Specifically, a wireless pressure sensor was threaded alongside the oesophagus and implanted into the thoracic cavity, where it could detect thoracic pressure changes that directly reflect respiration (Reisert et al., 2014). This approach was applied in awake, freely moving animals, allowing respiration to be monitored in a natural environment and during behavioural tasks.

In the current study, we present a new surgical approach to implant a wireless thoracic pressure sensor into the jugular vein, and verify the thoracic pressure signal against a head-fixed flow-sensor recording. Probe insertion into the jugular vein, rather than alongside the oesophagus wall (Reisert et al., 2014) simplified the postoperative care, with no requirement for a change to liquid-based diet. This surgical approach is thus more suitable for typical learning assays where water restriction is required for training. We then use this technique in conjunction with implanted EEG and EMG electrodes to measure brain activity in freely moving mice. We examined respiration pattern changes as mice voluntarily explored novel objects, novel odours and food odours, and also across different arousal states, while simultaneously examining the relationship between respiration and specific brain rhythms. Gaining insight into how respiration relates to brain activity and behaviour will provide new opportunities to understand how animals process the world around them and interact with their environment.

## RESULTS

### Implanting thoracic pressure sensor in the right jugular vein of mice

To measure respiration in freely moving mice, we implanted 6-8 week old mice with a wireless pressure sensor (Fig. 1A-F). Briefly, after exposing the right jugular vein, we ligated the superior portion of it to stop any blood flow (Fig. 1C). Next, we made a small incision (Fig. 1D) through which we inserted the pressure sensor towards the thoracic cavity while monitoring the pressure signal online (Fig. 1E). Once we reached the optimal position of the probe, the sensor was held in place with an additional ligature around the vein (Fig. 1F). (Figure 1; described in detail in the Materials and Methods section).

**Figure 1.**
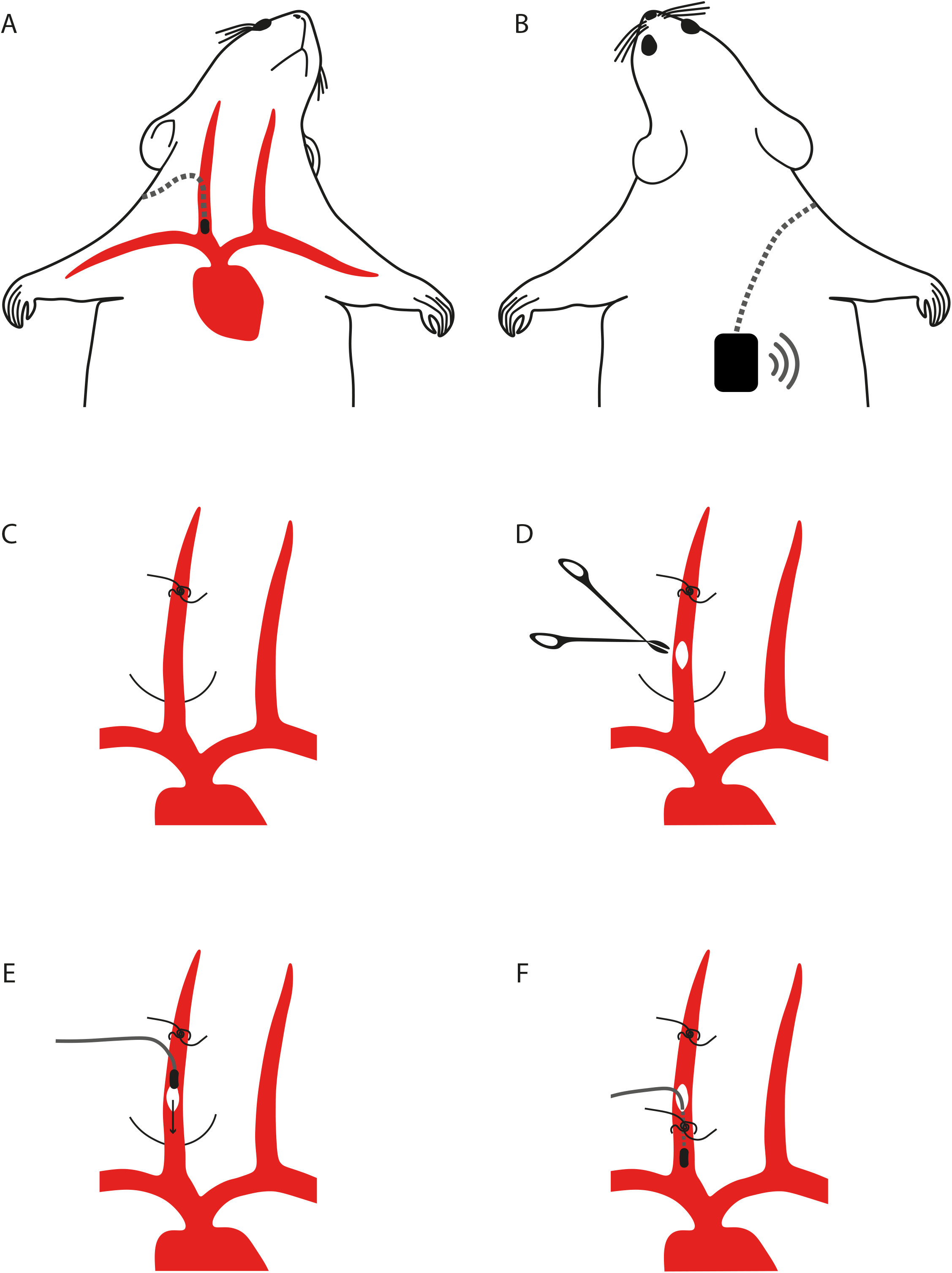
Jugular vein implantation of thoracic pressure sensor. (A) Schema showing the final location of the pressure sensor tip in the jugular vein and the subcutaneous location of the transmitter on the back (B). (C-F) The critical steps of the surgery are tying the rostral knot on the right jugular vein (C); making the incision on the jugular vein (D); inserting the sensor tip through the incision towards the heart (E) and securing the sensor at the correct position for optimal signal (F).

### Comparing thoracic pressure sensor and flow sensor signals

After recovering from the surgical implantation (> 7 days after surgery), mice were transferred onto a treadmill. After a short habituation session on the treadmill, we recorded the change in thoracic pressure signal using the implanted pressure sensor, and the change in nasal flow using a flow sensor placed in front of the nostrils (Fig. 2A). Both the sensors reflect respiration in the animals. Qualitatively, the signals looked similar, with downward deflection representing inhalation in both recording methods (Fig. 2B, Supp Fig. 2.1). To quantitatively compare these methods, we measured the frequency and peak inhalation time in both signals recorded simultaneously (Fig. 3). Using autocorrelation to estimate the average frequency, we did not observe any significant difference in the estimated frequency (R = 0.914; Linear regression) between the two signals (Fig. 3A-C) and the relative error between respiration frequency measurements derived from thoracic pressure and flow sensor measurements was 0.016±0.009 (mean±SD; n=21 traces; 4 mice, Fig. 3D).

**Figure 2.**
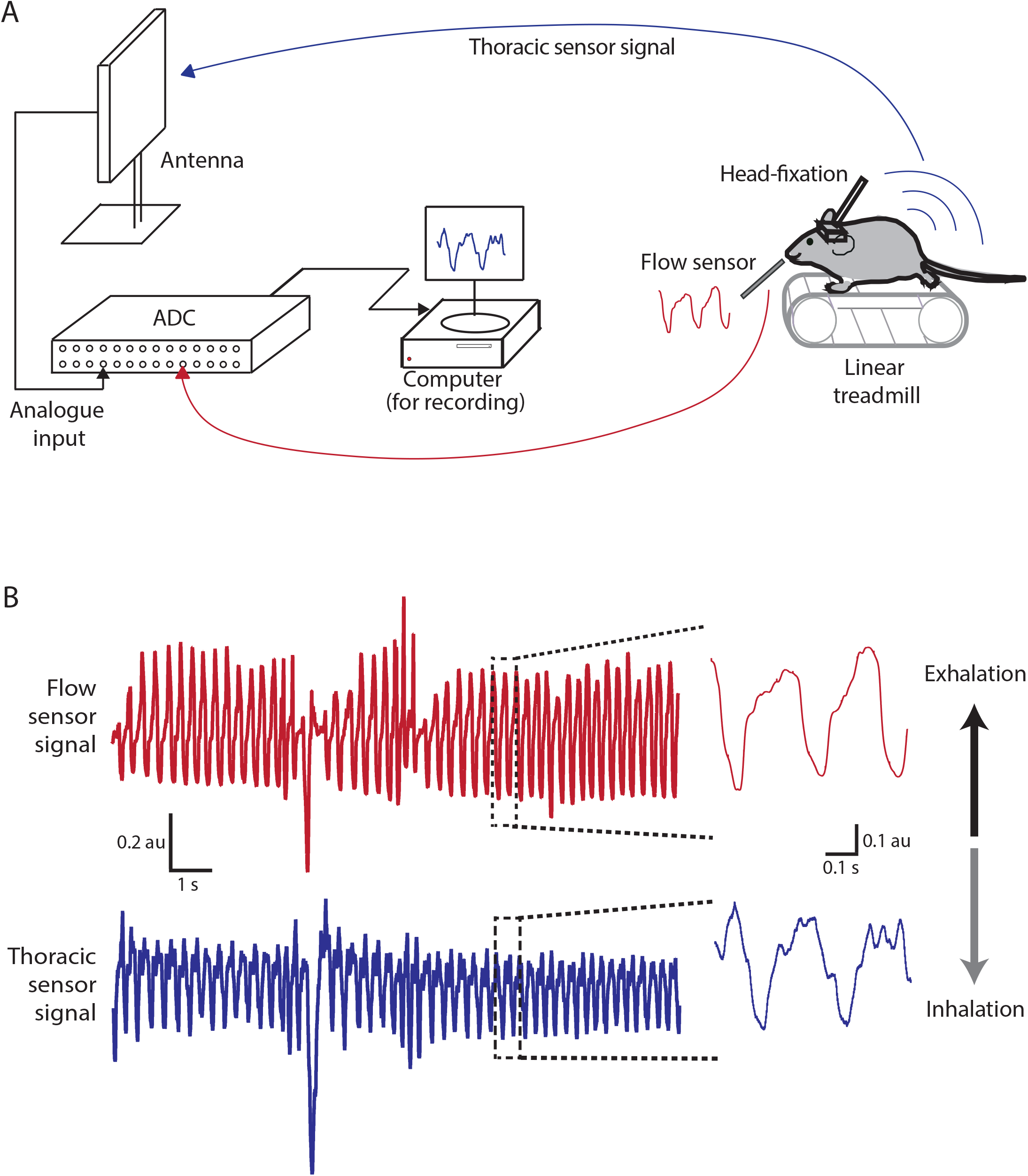
Head-Fixed experimental setup for simultaneous measurement from nasal flow sensor and thoracic pressure sensor. (A) A schematic diagram of the experimental setup used for simultaneous measurement of real-time respiration signals using a flow sensor placed near the nostrils and a thoracic sensor implanted in the jugular vein of head-fixed mice. (B) An example recording obtained simultaneously from the flow sensor (red) and the thoracic sensor (blue). A zoomed section is shown at the right.

**Figure 2.1.**
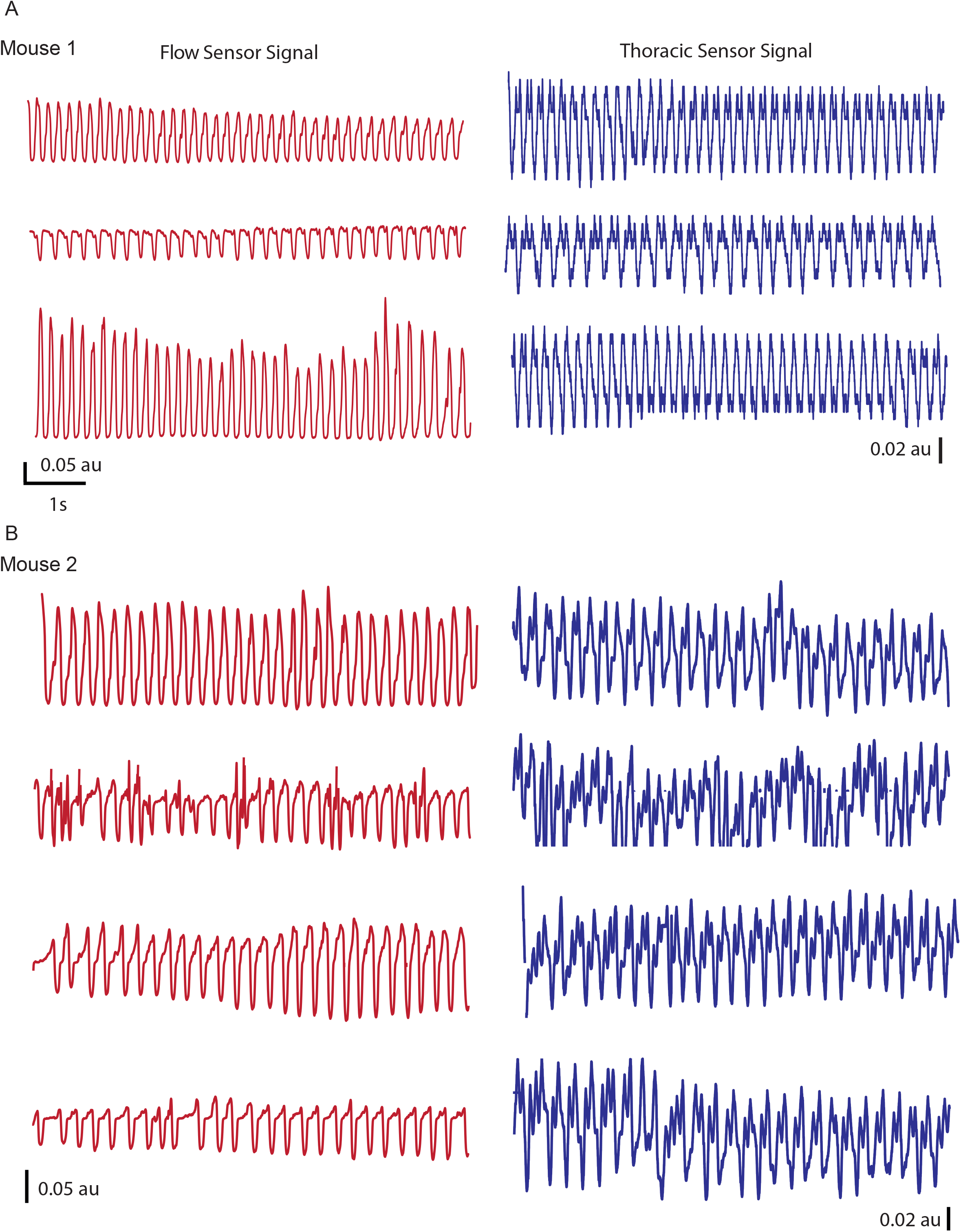
Example flow and thoracic recordings from two animals. Example recordings from two animals, obtained simultaneously from the flow sensor (left, red) and the thoracic sensor (right, blue). Top three traces (A) are from one mouse and bottom four traces are from the second mouse (B).

**Figure 3.**
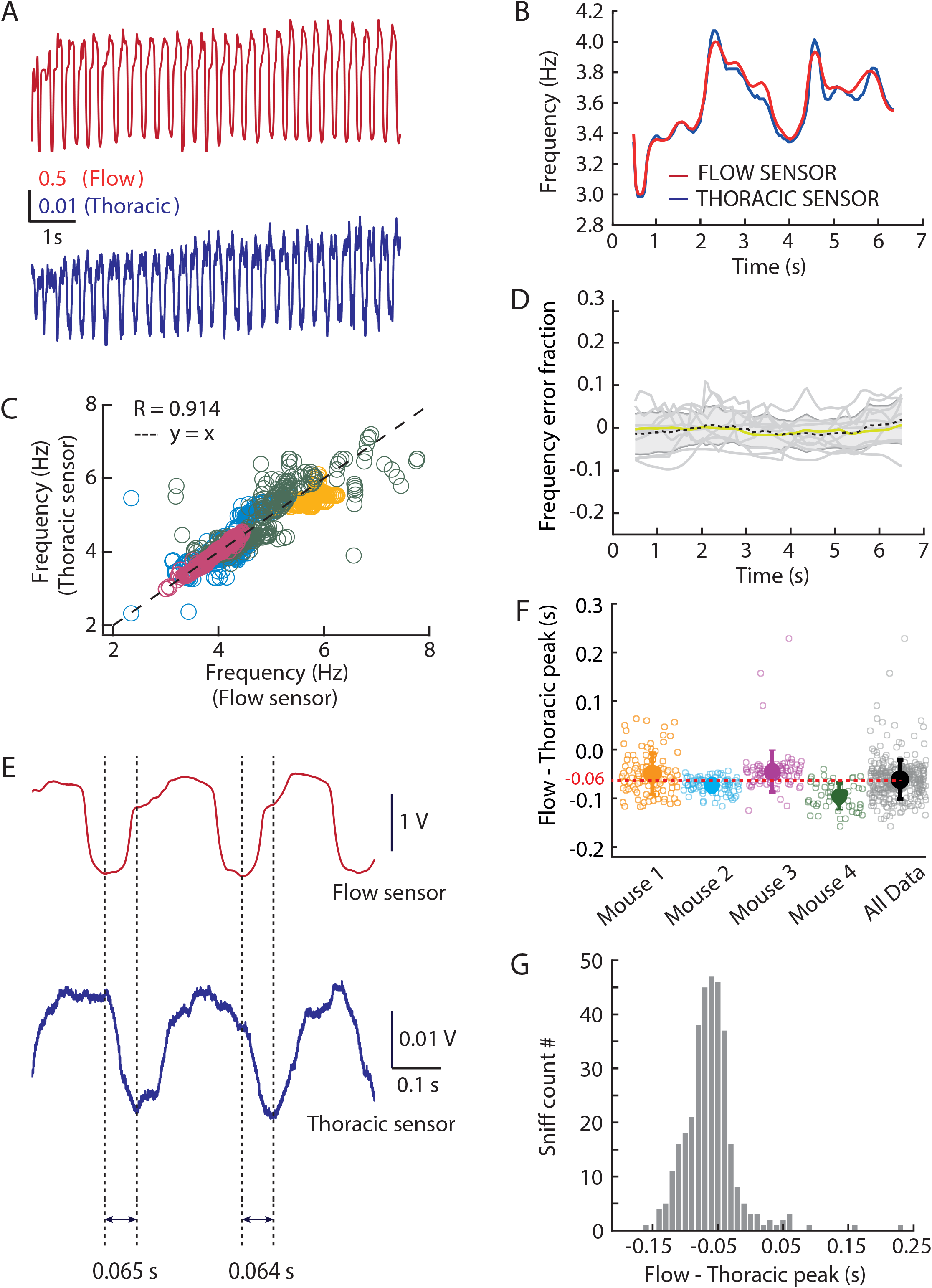
Implanted thoracic sensor is a reliable way of measuring respiration in awake mice. (A) An example recording obtained simultaneously using a flow sensor (red) and a thoracic sensor (blue) in a head-fixed mouse. (B) Average frequency (estimated using autocorrelation with a rolling 1 sec window, see Methods) from the 2 traces in (A). (C) Frequency estimate using the thoracic sensor vs flow sensor. Each marker represents the average frequency obtained from 1s segment of a trial. (Linear Regression analysis, R = 0.914, P<0.001). Different colours represent different animals. (D) Error fraction in estimating frequency from the same traces as in (C). (E) An example stretch of sniffs demonstrating the consistency of time-interval between the measured inhalation peak using the 2 sensors. (F) Time-interval between the inhalation peak times measured from the 2 sensors simultaneously. The colours represent the same animals as in (C). Solid markers represent mean±SD. The black marker represents the population average time-interval of -0.06 ± 0.03 s. (G) Histogram of the population of time difference in inhalation peak obtained from all the sniffs in (F), n=350.

As nasal flow and thoracic pressure reflect different aspects of respiration, there is likely to be a consistent shift in the time at which each signal peaks and knowing this shift could enable “translation” between these signals. We observed that the end of inhalation corresponds to the point of minimum thoracic pressure, and that the peak of inhalation preceded this point by 0.06±0.03 sec (n=350 sniffs; 4 mice). Overall, therefore, we have found that the thoracic pressure signal can be reliably used to measure respiration and to detect the time of peak flow inhalation. While we assume that this linear relationship holds for the higher frequencies observed during active exploration (below), it is an important caveat that we could only experimentally describe this relationship for the frequencies naturally demonstrated under head-fixation conditions (∼2-8 Hz). Interestingly, Reisert et al. (2014) were able to describe a similar relationship between thoracic pressure and intranasal pressure across a wider range of sniffing frequencies, and found that the variability in the difference between these two pressure signals was smaller at higher frequencies (their Figure 5).

### Freely moving mice have a bimodal respiration frequency not associated with running speed

Next, we used this method to measure respiration in freely moving mice while they explored an open arena. The thoracic pressure recording was acquired after a short period (∼2-3 mins) of habituation in a novel arena while movements were video-recorded for post analysis (Fig. 4A-B). The population of respiration recordings from all animals revealed a bimodal distribution of respiration frequency (Fig. 4C) with a small peak at approximately 3 Hz and a larger peak around 11 Hz. (Note that a bimodal distribution was also observed by Reisert et al., 2014, but in their study it was the lower frequencies that were more common. We suggest that this difference may come from the recording arenas – while the Reisert et al.’s mice were in their home cage, our mice were in a large, novel arena, which may have promoted more exploratory sniffing behaviour in the higher frequency range.) The estimated velocity from simultaneously acquired video recordings also showed a bimodal distribution (Fig. 4D) with peaks at approximately 7 cm/s and 40 cm/s. To understand whether respiration frequency was linked to running speed, we quantified the average velocity associated with 2 prominent frequency bands (1-5 Hz and 9-13 Hz) (Fig. 4E) and the average frequency associated with the 2 prominent bands of velocity (1-10 cm/s and 35-45 cm/s) (Fig. 4F). Surprisingly, we did not observe any significant difference in either of the cases (velocity comparison: p = 0.4691; and frequency comparison: p = 0.2501; both unpaired t-tests), suggesting that respiration rate was largely independent of running velocity (Correlation Co-efficient = -0.0071, Fig. 4G). While it is possible that some feature(s) of behaviour that we have not measured may predict respiration rate, these results suggest that respiration rate is not simply linked to motor activity, and fits with previous work showing similar differences in respiration rhythm (peaks at 3-5 Hz or 9-11 Hz) during head-fixed running on a treadmill (Nguyen Chi et al., 2016).

**Figure 4.**
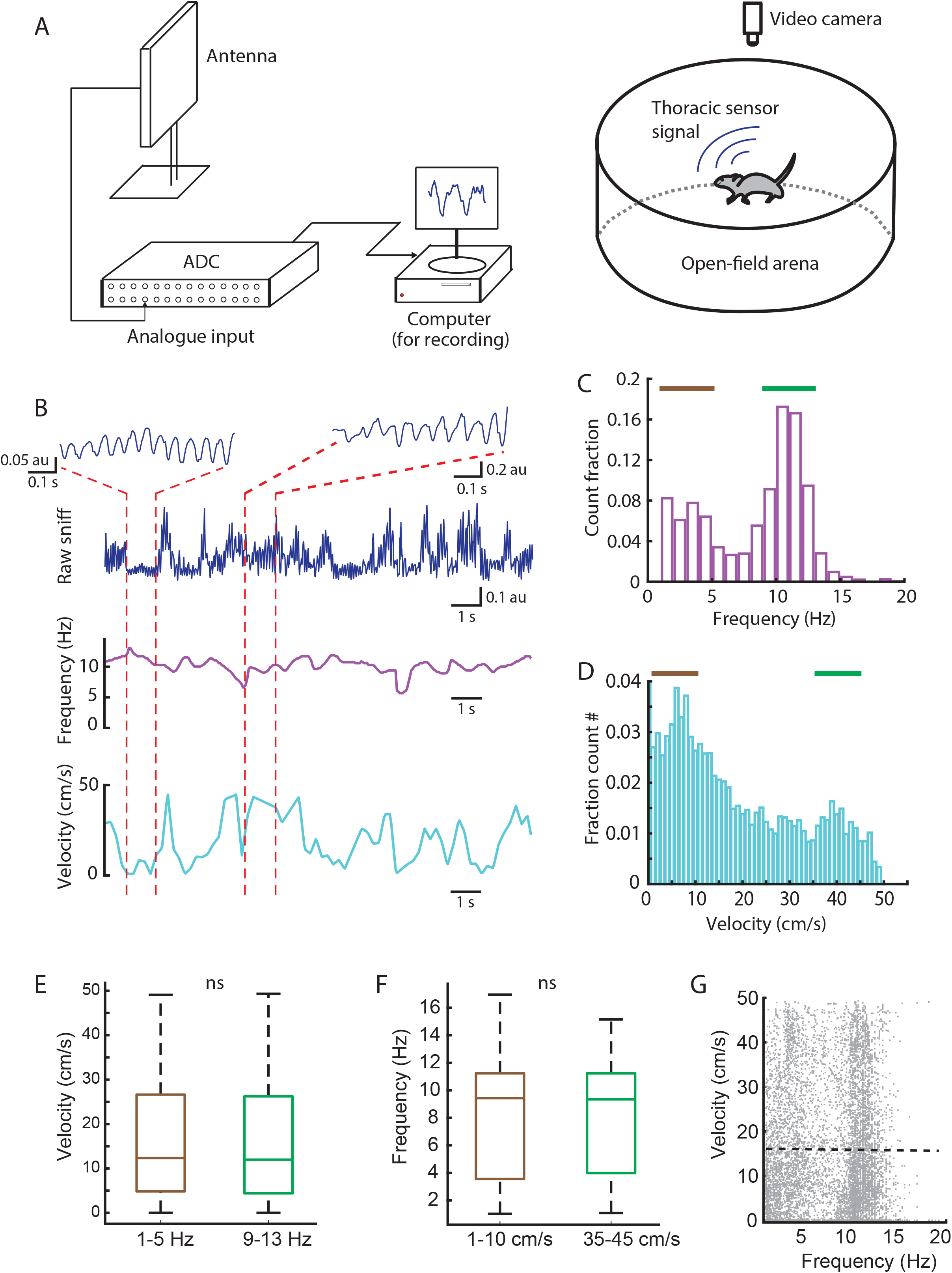
Respiration measurement from freely moving mice implanted with thoracic sensors exploring an open arena. (A) A schema of the experimental setup used for telemetry recording of respiration in freely moving animals in an open arena. (B) An example trace of respiration recording from an animal while it freely navigated in the open arena (*Top*). Note the respiration profiles in the 2 zoomed section during high frequency respiration, low velocity movement (*left*) and high frequency, high velocity (*right*). (C) Histogram of respiration frequency estimate from all animals (n=4 animals). Brown and green indicate the range of frequencies analysed in (F). (D) Histogram of running velocity from all animals in (C). Brown and green indicate the range of velocities analysed in (E). (E) Average running velocity was independent from respiration frequency in frequency ranges between 1-5 Hz (brown) and 9-13 Hz (green, indicated in C). (F) Average respiration frequency was independent from running velocity at speeds of 1-10 cm/s (brown) and 35-45 cm/s (green, indicated in D). (G) Running velocity vs. respiration frequency does not show any substantial correlation (R = -0.0071, dotted line) from all the animals in (C).

### Exploration of environmental cues significantly increases respiration frequency

We next asked how respiration frequency changes when mice encounter cues such as novel objects (e.g. rubber duck, plastic toys, empty bottles), monomolecular chemical odourants (ethyl butyrate, 2-Hexanone, eucalyptol and amyl acetate) and food (inaccessible pellet). After habituation to an empty arena, we placed a cue from each class into the arena, one at a time in a random order (Fig. 5A-C). We monitored respiration as the mice voluntarily explored these different cues (Fig. 5D-F). We then calculated the respiration frequency across all exploration bouts (Fig. 5G-I) and quantified the change in respiration frequency from the baseline after aligning the trials to the initial exploration video frame (Fig. 5J-L). For all the 3 classes of cues, we observed that the mice increased their respiration frequency significantly (food odour: p = 0.0103; novel object: p = 2.6e-5; and novel odour: p = 0.0394; paired t-tests) during exploration (Fig. 5M-O). Sniffing rates are well known to increase during tasks which rely on olfactory discrimination (Verhagen et al., 2007; Wesson et al., 2008a,b; Coronas-Samanos et al., 2016; Esquivelzeta Rabell et al., 2017; Jordan et al 2018a,b), but the present results, along with other results showing increased sniffing during exploratory locomotion even in the absence of any cues (Zhong et al., 2016), suggest that raised respiration may be a general feature of exploration. In turn, neural entrainment to this respiration rhythm may act similarly – but distinctly – to theta oscillations, providing a scaffold for long-range network communication across the brain (Nguyen Chi et al., 2016). We therefore sought to examine how the respiration changes during voluntary exploration are related to brain-wide activity in defined frequency bands.

**Figure 5.**
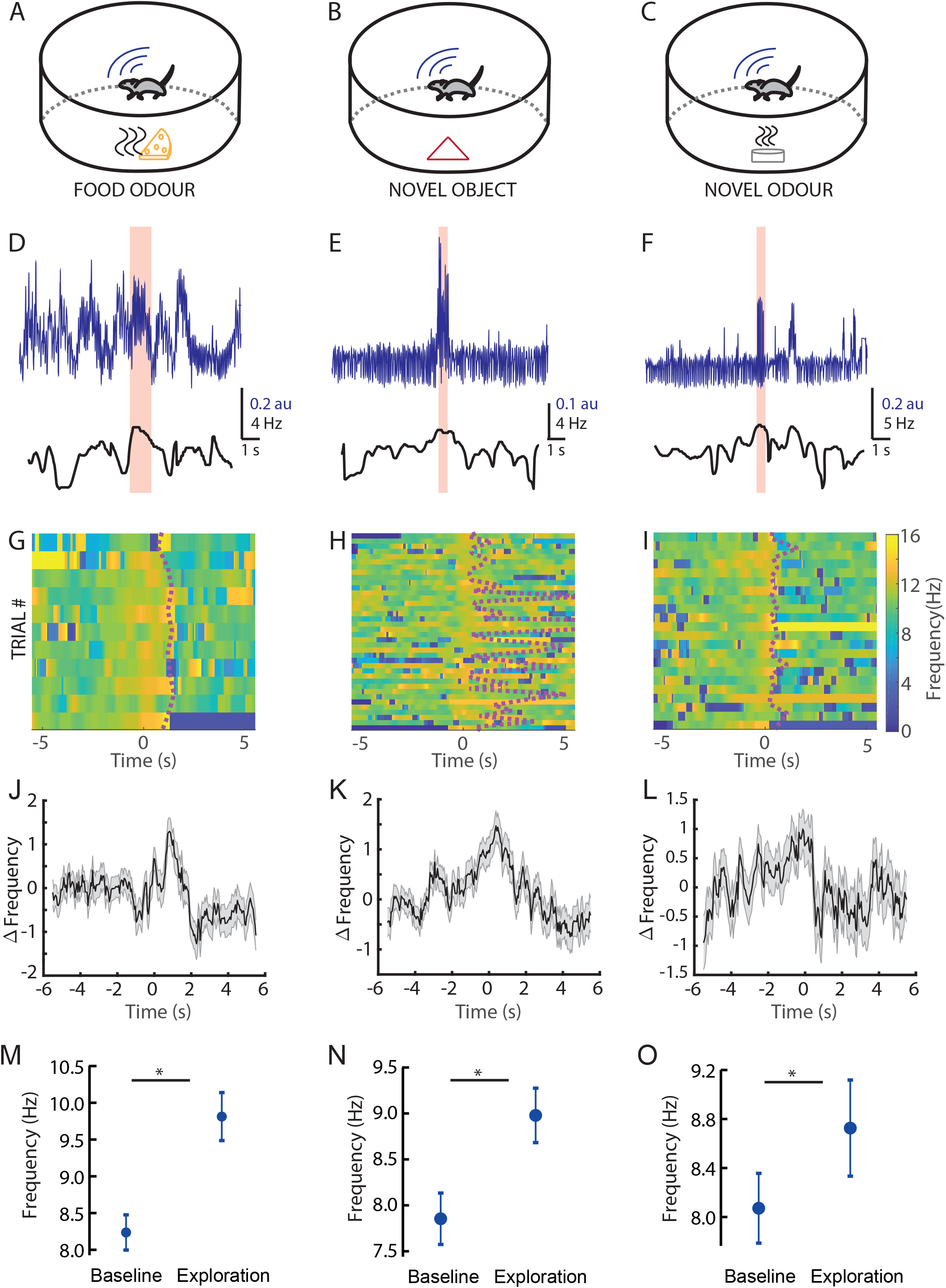
Mice show increased respiration frequency during exploring novel food, object and odour. (A) A schematic diagram of the experimental setup used for mice exploring food, (B) a novel object and (C) a novel odour while recording their respiration. (D) An example respiration recording (*top*) and its corresponding respiration frequency estimate (*bottom*) for a trial where an inaccessible piece of food was placed in the arena. The cream bar denotes the time of exploration in that trial (E) Similar example for a trial in which a novel object and novel odour (F) were placed in the arena. (G) Respiration frequency plots from multiple trials performed by an animal exploring the inaccessible food, novel object (H) and novel odour (I). Note the change in sniff frequency during the exploration time. Dotted line indicates the time at which the animal stopped exploring for each trial. (J) Baseline subtracted respiration frequency plotted against time for inaccessible food (n=85), novel object (n=134) (K) and novel odour (n=72) (L). The thick lines represent the mean and the shaded region represents the sem (4 mice). (M) Respiration frequency during the period of exploration significantly increases from the baseline respiration frequency for inaccessible food (p = 0.0014), novel object (p<0.001) (N) and novel odour (p = 0.03) (O) (4 mice).

### Exploration-triggered respiration changes increase or decrease the coupling between respiration and brain activity depending on EEG frequency band

Recent results have shown that cortical dynamics can be altered by changes in respiration, which were triggered experimentally by changes in CO_2_ levels (Girin et al., 2021). We sought to test how the sniffing changes during voluntary exploration relate to cortical dynamics.

Using the implanted thoracic pressure sensor in combination with simultaneous EEG/EMG recording (Fig. 6A-B), we explored how animals’ respiration related to brain activity during free exploration. In order to assess the relationship between EEG, EMG and respiration, we applied a frequency-domain locally-stationary time series analysis framework to the recorded data (see Methods). First, we assessed the EEG-EMG-respiration relationships in an open empty arena under the assumption of stationarity. This allowed us to build a baseline picture of the relation between all four recording channels (2xEEG, 1xEMG, 1xrespiration). We found that, in this state, respiration was directly related to both the EMG channel and the two EEG channels (which were also directly related to each other; Fig. 6C).

**Figure 6.**
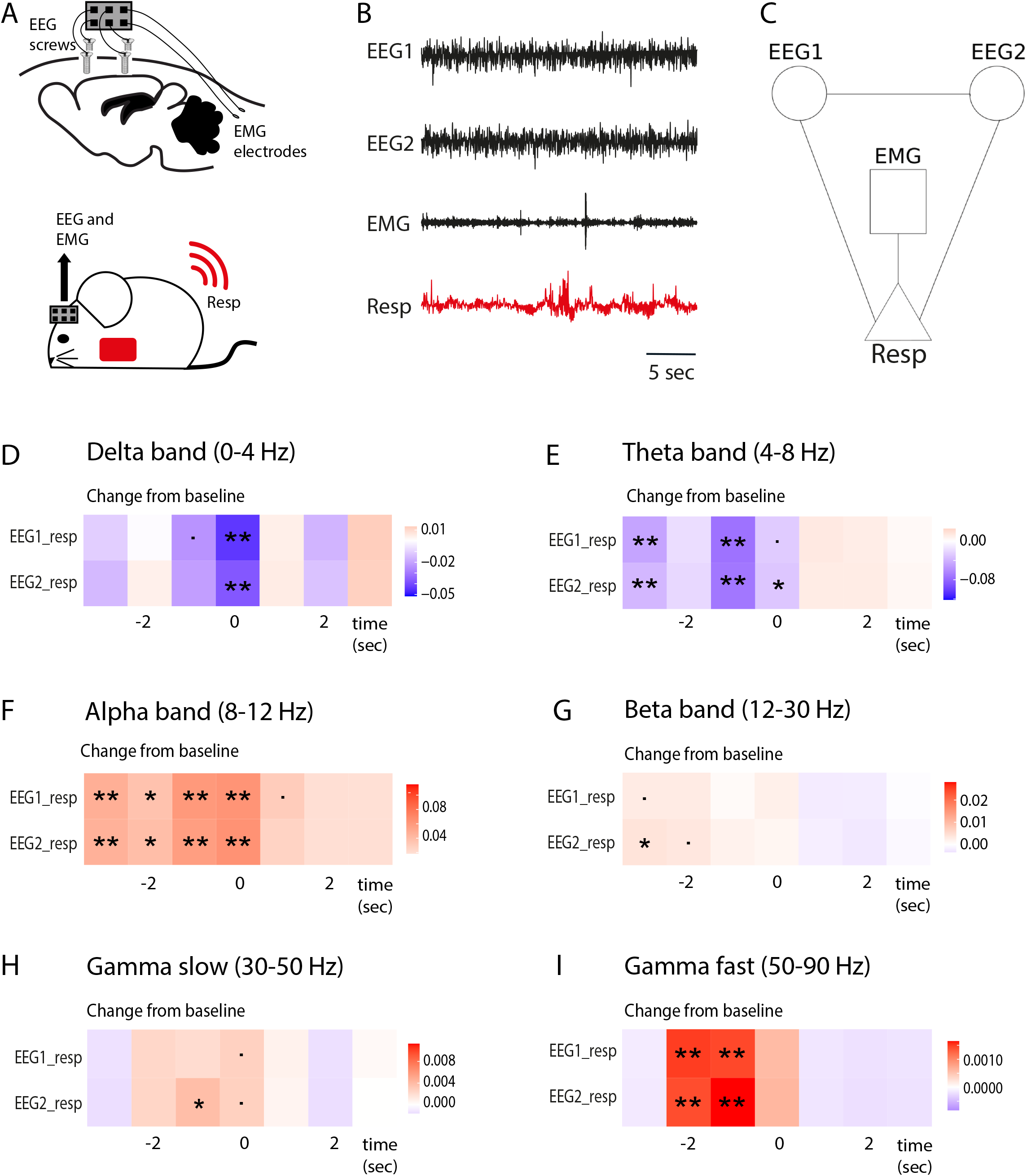
EEG, EMG and sniff relationships during freely moving awake behaviour. (A) Schematic of EEG/EMG electrode placement (top) and mouse implanted with both thoracic sensor and EEG/EMG recording devices (bottom). (B) Example raw traces from each of the four recording channels. (C) Baseline relationships between the four recording channels were estimated while the animals were awake in an open arena, in the absence of additional environmental cues. A frequency-domain time series analysis framework under the assumption of stationarity revealed that the two EEG channels were directly related to each other and to the EMG channel, and the EMG channel was directly related to the respiration channel. This plot represents partial coherence between EEG1, EEG2, EMG and TSE in the delta band in the open arena experiment, under assumption of stationarity, and a line was drawn between any 2 channels if both (a) partial correlation > 0.15 and (b) p-value for partial correlation being greater than 0 was < 0.05. (D-G) Estimates of changes in partial coherence between channel pairs in time bins before (negative time lag), during (0 time lag) and after (positive time lag) an animal explores an introduced environmental cue (either novel object, novel odour or food odour). Analysis was done separately for different frequency bands: delta (D; 4 mice), theta (E; 4 mice), alpha (F; 4 mice), beta (G; 4 mice), slow (H; 2 mice) and fast (I; 2 mice) gamma. Key for significance: ‘**’ = p<0.01; ‘*’ = p<0.05; ‘.’ = p<0.1.

We then used a frequency-domain approach called partial coherence to understand how the links between these channels evolve over a period of exploration. Partial coherence is a non-parametric measure of direct relationships between signals in a multivariate set. It is derived from the spectral density of the entire channels system, at any given frequency. It is valued between 0 and 1, and can be interpreted as the fraction of power shared by any 2 channels (in our case, EEG and respiration), while controlling for the influence of other channel (EMG) (Schneider-Luftman & Walden., 2016; and see Methods for more details).

For six frequency domains: (1) delta band, 0-4 Hz; (2) theta band, 4-8 Hz; (3) alpha band, 8-12 Hz; (4) beta band, 12-30 Hz; (5) gamma slow, 30-50 Hz; and (6) gamma fast, 50-90 Hz, we examined how the association between EEG and respiration channels changed before, during and after the mice explored environmental cues (whilst controlling for the influence of muscle activity recorded in the EMG channel), using a slowly-evolving locally-stationary (SeLV) framework. We found that, for the delta and theta frequency bands, the partial coherence across channels decreased relative to baseline before and during an exploration bout, increasing again after the animal stopped exploring (Fig. 6D-E). In contrast, for the alpha, beta and gamma frequency bands, the partial coherence across channels increased relative to baseline before and during exploration, decreasing again after the animal stopped exploring (Fig. 6F-I). This suggests that there is a direct relationship between respiration and brain activity, not mediated by muscle movement, which fluctuates significantly during exploration bouts. This change in direct signalling between brain and respiration differed based on frequency band: The relationship between respiration and brain activity in the delta and theta range decreased before and during exploration, while for the higher frequency bands – alpha, beta and gamma – the relationship between brain activity and respiration increased before and during exploration. For all frequency bands, these trends were independent of cue class (novel object / chemical odourant / inaccessible food). Interestingly, Zhong et al. (2017) also found that exploration was associated with increased alignment between respiration rhythm and fast gamma oscillations within the olfactory bulb and the prelimbic cortex, and our results suggest that this could be a more widespread pattern across the cortex. They also extend the picture across frequency domains, and it is interesting to note that the decreased relationships between respiration and brain activity occur in frequencies normally associated with rest or sleep (delta and theta), whereas increased association between respiration and brain activity are found within frequencies normally associated with arousal, sensory engagement and attention (alpha, beta and gamma).

### Respiration patterns vary across vigilance states

We next examined how respiration varied as the animal cycled through periods of wake and different sleep stages in its home cage. Similar to previous work using different methods of respiration recording (e.g. implanted thermocouples, Zhong et al., 2017; whole-body plethysmograph, Girin et al., 2021), we found that respiration patterns were different between different sleep states (Fig. 7A), tending to be high in frequency and amplitude but highly variable during wake; very regular and low in frequency and amplitude during NREM sleep, and slightly irregular in both frequency and amplitude during REM sleep. Switches between these different respiration signatures occurred almost instantaneously upon transition between different vigilance states (Fig. 7B). The distribution of inter-sniff intervals was different between sleep states, leading to a mean respiration frequency that was highest in wake (mean frequency = 6.79 Hz) and lowest in REM sleep (mean frequency = 3.09 Hz) (Fig. 7C).

**Figure 7.**
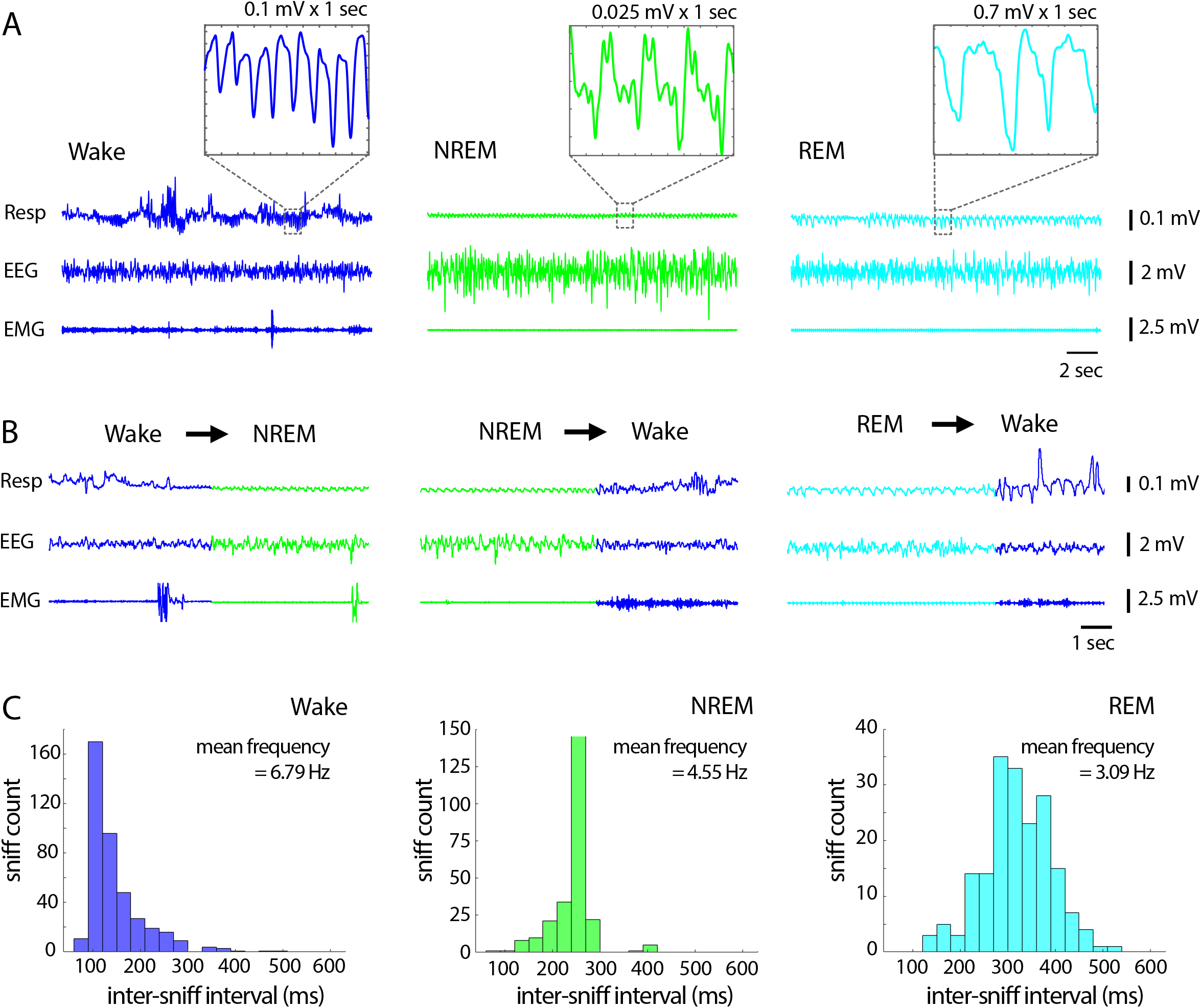
Respiration frequency changes with vigilance state. (A) Raw example sniff, EEG and EMG signals in wake (blue), NREM (green) and REM (cyan) sleep. The respiration trace (top row) shows that respiration is varied and often large in amplitude during wake, highly regular and low in amplitude during NREM sleep, and slightly irregular during REM sleep. Respiration frequency tends to be high during wake and lower in NREM and REM sleep (zoomed in boxes). (B) Changes in respiration are seen almost instantaneously upon transition between different vigilance states (note, mammals typically do not transition from REM to NREM sleep). (C) Distributions of inter-sniff intervals for wake, NREM and REM sleep, with mean respiration frequency for each state given in Hz.

### Odour-triggered respiration changes during NREM sleep do not induce changes in sleep-associated brain rhythms

Girin et al., (2021) found that the respiration changes induced by alterations in CO_2_-enriched air were capable of driving brain activity changes during wakefulness but not during sleep. We wanted to explore whether stimulus-induced sniff changes might tell a different story during sleep. We allowed the animal to rest in its home cage while monitoring EEG and EMG signals continuously (Fig. 8A). When the animal entered NREM sleep, we turned on the thoracic pressure sensor (it was not on continuously due to battery constraints – see Methods for details), and subsequently presented an odour to the sleeping animal (Fig. 8B). Within the 20 seconds following odour presentation, animals transitioned to wake 25% of the time, to REM sleep 25% of the time, and stayed in NREM sleep 50% of the time (Fig. 8C-D). When no odour was presented, the animal was more likely to stay in NREM sleep, and respiration did not change (Fig. 8D-E paired t-test, p = 0.16). But in the cases where odour presentation did not trigger a transition out of NREM sleep, we were able to monitor the effects on respiration and brain activity. In such cases, respiration increased (Fig. 8F paired t-test, p = 0.021; cf no change 20 seconds later: Fig. 8G paired t-test, p = 0.79), but no significant change was observed in sleep-related EEG rhythms (Fig. 8H-J delta: p = 0.40; theta: p = 0.31; theta:delta ratio: p = 0.91, paired t-tests). Thus, our results align with Girin et al.’s findings: neither CO_2_-induced nor stimulus-induced sniff changes appear to drive brain activity changes during sleep.

**Figure 8.**
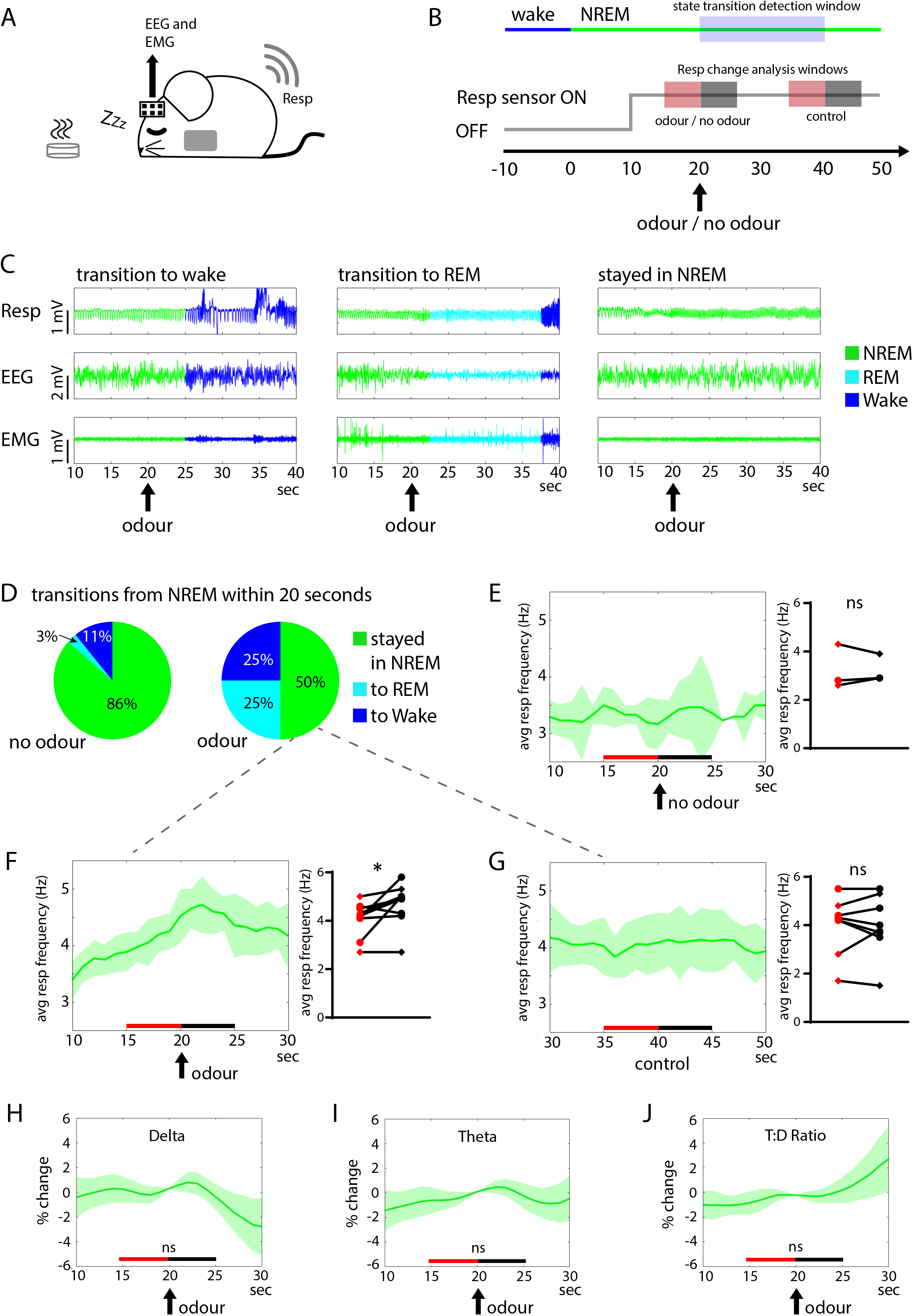
Odour presentation during sleep. (A) Schematic of recording set up. While the animal was sleeping, EEG/EMG and respiration signals were recorded, and odours were placed in the animal’s home cage. (B) Schematic of analysis framework. Ten seconds after the animal entered NREM sleep (switch from blue to green, top row), the respiration sensor was turned on (for one minute; grey line, middle row). Ten seconds later, odour was placed into the cage and the window for detecting state transitions was opened (for 20 seconds, blue shaded square). The red and black shaded squares represent the timing of respiration frequency comparisons (panels E-G) and EEG frequency band comparisons (H-J). (C) Raw example traces of respiration, EEG and EMG signals for an instance when the animal transitioned to wake, REM or NREM, within 20 seconds of odour presentation. (D) Pie charts showing proportion of state transitions that occurred between 20-40s from the onset of NREM sleep, for cases where odour was not presented (left pie chart) and cases where odour was presented at t=20s (right pie chart). While odour presentation did increase the proportion of transitions to wake and REM sleep, in half the cases, odour presentation did not cause the mouse to switch out of NREM sleep (10 out of 20 trials, across two mice). (E) In three trials, the respiration sensor was turned on without any odour being presented. In these cases, respiration frequency did not change 20 seconds after the start of NREM sleep (left: dark green line represents mean across trials, shaded region represents SEM; right: paired t-test, p = 0.16, two shapes represent two mice). (F) In the cases where odour presentation did not cause transition out of NREM sleep, there was a significant increase in the average respiration frequency (paired t-test, p = 0.021; colours and shapes as in E). (G) Control comparisons made 20 seconds after odour presentation (in cases where animal was still in NREM sleep) did not reveal a change in average respiration frequency (paired t-test, p = 0.79; colours and shapes as in E). (H-J) In cases where odour presentation did not cause transition out of NREM sleep, there was no significant change in the EEG delta power (H; p = 0.40), theta power (I; p = 0.31), or theta:delta ratio (J; p = 0.91) (colours as in E-G; p values are from paired t-tests, comparing average values for five seconds shown in red vs five seconds shown in black, as in E-G).

## DISCUSSION

We present a new implantation technique for a wireless thoracic pressure sensor, used to monitor respiration in conjunction with brain recording in freely moving mice. After calibrating with head-fixed respiration measurements using flow sensors, we were able to use this method to investigate respiration and its relationship to EEG brain activity in a variety of behavioural contexts: free exploration of novel environments and cues, and across different sleep-wake states.

While changes in respiration in response to olfactory discrimination odour cues have been well documented (Verhagen et al., 2007; Wesson et al., 2008a,b; Coronas-Samanos et al., 2016; Esquivelzeta Rabell et al., 2017), our data reveal comparable sniffing increases across voluntary exploration episodes of food and non-food odours, as well as novel objects. Along with other work showing increased sniffing during exploratory behaviours in absence of any specific cues, this suggests that raised respiration may be a general feature of active exploration, even for non-olfactory stimuli, and fits with observations that diverse sensory stimuli can arouse sniffing (Welker, 1964).

Given the increasing view that respiration may provide the scaffold for a brain-wide rhythm that can help coordinate neural information transfer across distant brain regions (Heck et al., 2017; Tort et al., 2018), we sought to understand how exploratory sniffing in a voluntary, freely moving task was related to brain activity in specific frequency bands. Using cortical EEG recording we found that, during stimulus approach, there was a decrease in the association between sniffing and both delta and theta frequency. Cortical delta frequency is most prominent during deep sleep, while cortical theta frequency is associated with REM sleep (reviewed in Poe et al., 2010), and so it is perhaps intuitive that the respiratory link with these rhythms would be reduced as the mouse makes an active exploratory approach. By the same token, it makes sense that it is the higher frequency domains (alpha, beta, gamma) that show increased partial coherence, as these rhythms are typically associated with arousal and attention (reviewed in Wang, 2010). The particularly strong increase in partial coherence with the fast gamma rhythm is similar to the results of Zhong et al. (2017), who found that – even in the absence of any specific cues – exploratory sniffing was associated with increased alignment between the respiratory rhythm and fast gamma oscillations in the olfactory bulb and prelimbic cortex. In general, our data show that active exploration of sensory stimuli in a voluntary, freely moving setting trigger sniff changes that show transiently increased association with specific brain activity rhythms.

Several recent studies have revealed that neuronal entrainment of different brain rhythms to the respiration rhythm is affected by arousal (Zhong et al., 2017; Cavelli et al., 2020; Girin et al., 2021; Tort et al., 2021; Karalis & Sirota, 2022), so we next looked at respiration across different sleep and wake states. We found that sniff amplitude, frequency and variability are different between wakefulness, NREM sleep and REM sleep, and that transitions between these vigilance states trigger instantaneous changes in sniff pattern. These results show that changes in brain state are linked to observable changes in sniff behaviour, even in the absence of any change in the olfactory environment, and are in line with what has previously been reported in human sleep (Reed & Kleitman, 1926; Gutierrez et al., 2016). Interestingly, odour introduction during deep sleep is less likely than other sensory stimuli to produce EEG signs of arousal in humans (Carskadon & Herz, 2004), and may even deepen sleep (Perl et al., 2016), although these studies did not examine respiratory changes. Here, we found that introducing an odour to the animal’s home cage during NREM sleep could trigger an increase in sniffing even in the absence of any vigilance state changes. This suggests that, during sleep, active sampling can be modulated even without any obvious changes to the overall arousal state of the brain. Our results also expand on those of Girin et al. (2021), who showed that breathing changes triggered not by an olfactory stimulus but by changes in CO_2_ concentration, were least capable of driving respiration-related brain activity changes during deep sleep.

In summary, we have used a new surgical technique to implant a telemetry-based thoracic pressure sensor, and have shown that this can be used to accurately measure respiration in freely moving mice. Since respiration rhythm is increasingly being viewed as an important brain-wide scaffold to which other neural rhythms can align, we combined this new measurement technique with implanted EEG and EMG to monitor brain activity in specific frequency bands during behaviour and across vigilance states. During stimulus exploration, the association between respiration and cortical delta and theta decreased, but the association between respiration with alpha, beta and gamma, increased. Odour presentation during sleep was able to cause a transient increase in sniffing, but did not appear to change dominant sleep rhythms in the cortex. Overall, our data align with the idea that respiration may be a useful driver for synchronising specific brain rhythms, particularly during wakefulness and exploration, but during sleep, respiratory changes seem less able to impact brain activity.

Having demonstrated the flexibility with which thoracic pressure sensing can be combined with different behavioural assays and brain recording, we anticipate that this wireless technique to measure respiration will provide many new insights into the way that animals use olfactory information to understand the environment. Our recent findings show that mice can compute sub-sniff level information (Ackels et al., 2021; Dasgupta et al., 2022). Using the present technique it will be possible to interrogate brain activities related to sub-sniff level computation in freely moving animals.

## MATERIALS AND METHODS

### Implantation of thoracic pressure sensor in right jugular vein

All animal procedures performed in this study were regulated and approved by the Institutional Animal Welfare Ethical Review Panel and the UK government (Home Office) under license PA2F6DA12. All the surgeries and behavioural assays were performed on C57/Bl6 mice.

Thoracic pressure sensors (Stellar implantable transmitter device, 10X normal gain, E-430001-IMP-22, TSE systems, Germany) were implanted in the right jugular vein in mice. 6-8 weeks old animals were put in individual cages 2 days before surgery to ensure acclimatization and proper intake of drugs orally. On the day prior to the surgery, 0.2ml of egg custard (Cow & Gate) + 0.2ml oral Metacam suspension was given in addition to freely available food. On the day of surgery, the animal was weighed and anesthetized using Fentanyl (0.05mg/kg) + Midazolam (5mg/kg) + Medetomidine (0.5mg/kg), delivered ip. Further, for analgesia Meloxicam (10mg/kg) + Buprenorphine (0.1mg/kg) was administered subcutaneously. The animals were then placed on a heatpad (DC Temperature Controller, FHC, USA) controlled by a rectally inserted temperature sensor. Body temperature was continuously monitored and maintained at 37 ± 0.5 °C. The probes consisting of two parts, a transmitter (2cm X 1cm X 0.3cm) and a catheter (5cm long and ∼0.4mm diameter), were implanted using aseptic surgery techniques. The skin on the right of the neck’s midline was shaved and disinfected with 25% (v/v) chlorhexidine. Next, a ∼2 cm skin incision was made and using blunt tools a subcutaneous tunnel was created underneath the right arm up to the back of the animal. Pre-sterilised saline solution was used to irrigate the wound regularly. The transmitter was pushed through this tunnel up to the back of the animal (Fig. 1B) while keeping the sensor end out of the wound. Next, post isolation of the right jugular vein, a knot was tied on the dorsal-most part of the isolated section of the vein using non-soluble surgical sutures (Fig. 1C). A small incision was made on the vein surface to insert the sensor tip of the catheter (Fig. 1D). During the process of insertion of the sensor tip (Fig. 1E), the pressure signal was continuously monitored to identify the best spot for placement. Upon reaching the best spot, a knot was firmly tied around the vein enclosing the catheter (Fig. 1F). The remaining suture thread from the dorsal knot was also used to make a knot around the catheter for extra stability of the placed sensor. Finally, the wound was closed using 6-0 silk suture and a reverse– cutting needle. The animal was recovered in a heated chamber after injecting 1.2mg/kg Naloxone + 0.5mg/kg Flumazenil + 2.5mg/kg Atipamezole (ip). Sterile saline (0.2 ml) was injected subcutaneously for faster recovery. The animals were monitored and their bodyweights were recorded regularly at least for the next 10 days post-surgery.

For the head-fixed recordings, a subset of animals also underwent a head-fixation implant attachment in the same surgery. Briefly, a custom-made head-fixation implant was glued to the skull with a medical grade superglue (Vetbond, 3M, Maplewood MN, USA). Further, dental cement (Palladur, Heraeus Kulzer GmbH, Hanau, Germany) was applied around the base of the implant to strengthen the fixation.

Post-surgery the animals were allowed at least a week to recover from the surgery and to get back to their pre-surgery body weights before being used for experiments.

### EEG and EMG electrode implantation

For the EEG/EMG exploration and sleep experiments, mice underwent a second surgery at least one week after implantation of the telemetry device. Mice were anaesthetized with isoflurane and injected s.c. with meloxicam (2 mg/kg of body weight) for analgesia. After positioning in a stereotaxic frame (Kopf Instruments), mice were implanted with four miniature screw electrodes (from bregma: AP +1.5 and ML +1.5 (ground); AP +1.5 and ML -1.5 (common reference); AP -1.5 and ML -1.5 (EEG 1); AP -1.5 and ML +1.5 (EEG 2) and two EMG electrodes (inserted into neck musculature). These electrodes were each connected, via an insulated wire, to a different gold pin of a EEG/EMG headstage. The EEG/EMG headstage was affixed to the skull using dental adhesive resin cement (Super-Bond C&B). Mice were allowed to recover for a further week before participating in head-fixed and then freely moving behaviour experiments.

### Data acquisition of the telemetric signal

A commercial telemetry system associated with the probes (TSE systems) was used for the wireless recording of the thoracic pressure signals in awake animals. The signal from the probe’s transmitter was sensed by the antenna of the telemetry system output which was connected to DAQ (CED Micro1401 with ADC12 expansion, Cambridge Electronic Design Limited, UK) and controlled by spike2 (Cambridge Electronic Design Limited, UK) on a computer. The signal was sampled at 1 KHz by the sensor and eventually digitized at 10 KHz.

Because the implanted probe has a fixed battery life, we could not acquire the thoracic pressure signal continuously. We therefore acquired in 0.5 – 2 minute bursts, with onset timed according to each experiment (before object introduction in the exploration experiment, and after online detection of NREM sleep in the sleep experiments).

### Data acquisition of the nasal flow sensor signal

A mass flow sensor (FBAM200DU, SensorTechnics) was used to measure the flow change in front of the nostril thus generating a continuous respiration signal as described previously (Dasgupta et al., 2022). The signal was digitized at 10 KHz simultaneously with the thoracic pressure signal using the same CED Micro1401 DAQ.

### Data acquisition of the EEG/EMG signals

EEG and EMG signals were recorded using the Pinnacle 3-channel tethered system (8200-K1-SL; Pinnacle Technology Inc). Signals were filtered by the preamplifier (high pass above 0.5 Hz for EEG and above 10 Hz for EMG) and then recorded in Spike2, via the same CED Micro1401 DAQ.

### Head-fixed experiments

The animals were placed in a custom-made head-fixation apparatus attached to a treadmill. The flow sensor was placed in front of the nostrils. Following a brief period of habituation (∼ 15 minutes), signals from the flow sensor and the thoracic sensor were recorded simultaneously. The signals were as described above.

### Behavioural experiments

During the behavioural sessions, mice were placed on the floor of a circular open-topped enclosure (50 cm in diameter) and video was recorded using a Raspberry Pi camera mounted on the ceiling (approximately 1.5 m above the enclosure). The camera sent out TTL pulses on a frame-by-frame basis (10 Hz) to the DAQ, which synchronized the respiration recording with the video recording. Formal animal tracking of head, centre and tail coordinates was performed offline using Ethovision XT software (Noldus). This allowed us to define exploratory approach and retraction times, and instantaneous velocity for all the frames of each trial. Four types of behavioural experiment were performed, while video, respiration signal and EEG/EMG (in a subset of mice) were recorded:

1. Open arena exploration: The mouse was gently placed in the arena and allowed to explore freely for approximately five minutes.
2. Novel object: Multiple objects (rubber duck, water bottle, nail varnish bottle, empty food hopper, black bottle, muffin toy) were placed in the arena one at a time, far away from the mouse, who was then allowed to approach and explore the object freely for approximately five minutes.
3. Novel odour: A glass petri dish containing a tissue piece impregnated with 2 ml of pure odour (ethyl butyrate, 2-hexanone, isopentyl acetate or eucalyptol) was placed in the arena, far away from the mouse. The mouse was then allowed to approach and explore the petri dish freely for approximately five minutes.
4. Inaccessible food: A food pellet was placed in a meshed container, which was then placed in the arena, far away from the mouse. The mouse was then allowed to approach and explore the container freely for approximately five minutes.

### Sleep experiments

EEG/EMG and respiration signals were recorded as the animal rested in its home cage, during the first half of the light phase. EEG signals were monitored online, to assess the arousal state of the animal in real time. When the animal entered NREM sleep, we waited for approximately ten seconds before turning on the telemetry respiration sensor for one minute. If the animal was still in NREM approximately ten seconds after that, we carefully placed a petri dish containing a tissue piece impregnated with 2 ml of pure odour (ethyl butyrate, 2-hexanone, isopentyl acetate or eucalyptol; as in the awake behaviour experiments), into the cage (as we did not see any consistent difference between odours, data was pooled across all four odours). Trials where the placement itself woke the animal up were excluded from analysis. The petri dish was left in position in the cage until sniff monitoring for that trial ended (one minute). Trials were separated by at least five minutes.

### Data analysis - Respiratory signals

The data was analysed using custom scripts written in MATLAB R2019 (Mathworks).

### Frequency estimate

The raw data was standardized and detrended. Next, one second of data were passed through an autocorrelation routine. The corresponding time of the first peak was estimated and its reciprocal was used to estimate the dominant frequency for that 1s period. This was repeated using a rolling window of one second with a shift of 50 ms between adjacent windows.

For the freely moving experiment with novel cues, we noted the frames of start and end of each exploration bout and included three seconds preceding and following each exploration bout to make up an individual trial. We followed the same method for estimating frequency for all the trials thus extracted. Events with baseline time less than three seconds were discarded. All trials were aligned to the start of exploration. We calculated the average baseline frequency from the first 500 ms of the trial and the average exploration frequency from the first one second of object exploration. We then subtracted the baseline frequency from the entire trial to plot change in frequency, and this was averaged over trials and across mice.

Frequency error calculation (Head-fixed): The frequency estimation for a 1s period was estimated from both the flow sensor data and the thoracic sensor data. Next, the relative error for that 1s was calculated as;

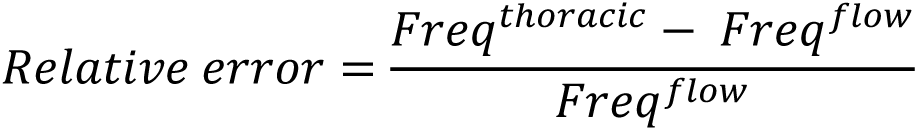

The relative error is then averaged over the entire trial period to get an average relative error for the specific trial. The mean and standard deviation is estimated from all the trials from all the animals.

### Inhalation peak detection

The peak of inhalation was detected as the trough of every sniff cycle in the flow sensor signal and the simultaneously recorded thoracic sensor signal. The peaks were detected using custom written scripts in MATLAB R2019 (Mathworks). The peak detection was manually scrutinized for discrepancies. To compare peak times between the thoracic and the flow sensor; inhalation peaks were detected in the signals recorded from both the sensors. Next, we calculated the difference in time between the 2 near simultaneous occurring peaks. Since the pressure change in the thoracic cavity happens slightly after any flow through the nostrils, it was assumed that the inhalation peak estimated from the flow sensor signal will precede that estimated from the thoracic sensor signal.

### Data analysis - Behaviour

Nose, centre, and tail tracking were performed using Ethovision software. We estimated mouse position, head direction, and instantaneous velocity for each video frame of each trial. Mice were considered to have initiated an exploration bout as soon as their nose entered a 2 cm perimeter around the object/odour/food. Retraction time was defined as the next frame in which the mouse’s nose moved out of this perimeter space. In general, these definitions were not ambiguous, and classifying exploratory initiation and retraction times either automatically (using Ethovision) or manually (by scrolling frame-by-frame through the video) gave almost identical results.

### Data analysis - Vigilance state classification

Arousal states – NREM, REM and wake – were automatically classified using sleep analysis software in Spike2, and then manually verified in 5 second epochs (as in Harris et al., 2022). Wakefulness was defined as de-synchronised, low amplitude EEG and tonic EMG with bursts of movement. NREM sleep was defined as synchronized, high amplitude EEG in the delta frequency range (1-4 Hz) and reduced EMG activity relative to wakefulness. REM sleep was defined when EEG had reduced delta power but prominent power in the theta range (4-10 Hz), and EMG showed an absence of muscle tone.

### Data analysis – Association between EEG, EMG and novel Respiration (Resp) signals

Data recorded from the 4 physiology channels (EEG1, EEG2, EMG and Resp) during each experiment were pre-processed before analysis. First, the EMG and Resp channels were downsampled to the sampling rate of the EEG channels (400 Hz). Exploration behaviour markers derived from video recording were then aligned with the timeline of the physiology recordings, using TTL time frame alignments. To save battery, the respiratory sensor was not activated throughout the whole experiment, and its recording was interrupted during some experiments. Thus, we removed all data points which had no Resp recording. The resulting data was then band-pass filtered with a Butterworth filter, with band pass 0.5-120 Hz.

For each experiment, the resulting dataset was a 4-dimensional time series, which was then analysed in six frequency bands of interest: Delta (0.5-4 Hz), Theta (4-8 Hz), Alpha (8-12 Hz), Beta (12.5-30 Hz), Gamma slow (30-50 Hz) and Gamma fast (50-90 Hz). For each frequency band and each experiment, the data was filtered again using a low-pass / band pass filter, with appropriate filter settings.

We then applied the slowly evolving locally stationary process (SEv-LSP) framework from Fiecas & Ombao (2016) to all exploration experiments. The four channels [EEG1,EEG2,EMG,Resp] recorded during each experiment were treated as a non-stationary 4-dimensional time series. The 4-dimensional time series were cut into small epochs (0.5-2 sec long) which were treated as “locally stationary”. The choice of epoch length was chosen differently for each frequency band, to ensure that a single bin always contained approximately ten full cycles: Delta band: 2.22 sec; Theta band: 0.83 sec; Alpha band: 0.5 sec; Beta band: 0.23 sec; Gamma slow: 0.125 sec; and Gamma fast: 0.07 sec.

The open arena experiments, which included no exploration bouts, were analysed using a stationary framework, meaning that it was not cut into epochs but analysed as one time-block. The rest of the analysis was otherwise identical to the one applied to all other experiments.

For each experiment, and for each time epoch, the spectral power matrix was estimated using Thomson’s multitaper estimate (Riedel & Sidorenko, 1995) with K=p x 1.5 = 6 sine tapers. After regularisation (Schneider-Luftman, 2016; Schneider-Luftman & Walden, 2016), the spectral matrix estimates were used to derive partial coherences. The partial coherence is a non-parametric measure of conditional independence between pairs of time series, and was used here to assess the relationship between the [EEG1,EEG2,Resp] channels inside each epoch. With p=3 channels, we have Npcoh=3 relationships between pairs of channels. Partial coherence estimates and p-values (H0: partial coherence =0) were derived for all 3 pairs of channels, at each epoch. Results were adjusted for false discovery rate (FDR) across number of pairs and number of epochs in the experiment. This analysis framework produces a set of slowly evolving partial coherences between the [EEG1,EEG2,Resp] channels, for each experiment and each frequency band of interest, and produces a set of epoch-sized partial coherences for each pair of channels.

This analysis framework applied to the open arena recordings produced one set of partial coherence estimates, which can be represented in a graphical model (Fig. 6C). This plot represents partial coherence between EEG1, EEG2, EMG and TSE in the delta band in the open arena experiment, under assumption of stationarity, and a line was drawn between any 2 channels if both (a) partial correlation > 0.15 and (b) p-value for partial correlation being greater than 0 was < 0.05.

For all other experiments, we then investigated the association between exploration bouts and evolutionary partial coherences between channels derived from the analysis framework described above. We pooled all experiments together and regressed partial coherence estimates against lagged exploration bouts (+/- 3 seconds around exploration bouts), using multiple-multivariate regression in R (version 3.6.1). This was done first on all types of exploration cues (novel objects, odour, food), then split by exploration cue type. The results of interest from the regression models are the estimated change from baseline in partial coherence values before/during/after exploration bouts, with the null hypothesis being that partial coherence values do not change around exploration bouts. This process was performed separately for each frequency band of interest.

## ACKNOWLEDGEMENTS

This work was supported by the Francis Crick Institute which receives its core funding from Cancer Research UK (CC2036), the UK Medical Research Council (CC2036), and the Wellcome Trust (CC2036); by the UK Medical Research Council (grant reference MC_UP_1202/5); a Wellcome Trust Investigator grant to ATS (110174/Z/15/Z) and the National Science Foundation/Canadian Institute of Health Research/German Research Foundation/Fonds de Recherche du Québec/UK Research and Innovation–Medical Research Council Next Generation Networks for Neuroscience Program (Award No. 2014217). We thank Yolanda Saavedra Torres and Rene Remie for helping in the standardisation process of the surgical procedure. We thank the animal facilities at National Institute for Medical Research and the Francis Crick Institute for animal care and technical assistance. We thank Matthew Wachowiak, Johannes Reisert and Mihaly Kollo for their helpful comments on earlier versions of the manuscript. For the purpose of Open Access, the author has applied a CC BY public copyright licence to any Author Accepted Manuscript version arising from this submission.

